# A Single Mutation in an Enteric Virus Alters Tropism and Sensitivity to Microbiota

**DOI:** 10.1101/2024.08.16.608258

**Authors:** Andrea K. Erickson, Danica M. Sutherland, Olivia L. Welsh, Robert W. Maples, Terence S. Dermody, Julie K. Pfeiffer

## Abstract

Many enteric viruses benefit from the microbiota, and depletion of the microbiota reduces infection of noroviruses and picornaviruses in mice. However, *Reovirales* viruses are outliers among enteric viruses. Rotavirus infection is inhibited by bacteria, and we determined that several reovirus strains have enhanced replication following microbiota depletion. We found that an isogenic pair of reoviruses have opposing infection outcomes after microbiota depletion. Microbiota depletion reduces infection of reovirus strain T3SA+ but increases infection of strain T3SA−. These strains differ by a single amino acid polymorphism in the σ1 attachment protein, which confers sialic acid binding to T3SA+. Sialic acid binding facilitates T3SA+ infection of intestinal endothelial cells, while T3SA− inefficiently infects intestinal epithelial cells due to restriction by microbiota-driven, host-derived type III interferon responses. This study enhances an understanding of the complex interactions of enteric viruses, the microbiota, intestinal tropism, and antiviral responses.

**Highlights:**

- Microbiota enhance infection of a sialic acid (SA)-binding reovirus strain but inhibit infection of an isogenic non-SA-binding reovirus strain that differs by a single amino acid change.
- SA binding facilitates viral infection of intestinal endothelial cells in a microbiota-dependent manner.
- A non-SA-binding reovirus strain inefficiently infects intestinal epithelial cells due to microbiota-driven type III interferon responses.

## INTRODUCTION

The digestive tract contains a remarkable microbial community, which influences mammalian virus infection by several mechanisms. For example, non-enteric viruses such as arboviruses, influenza virus, and lymphocytic choriomeningitis virus have increased replication in microbiota-depleted mice, which is likely due to bacteria-mediated priming of interferon (IFN) antiviral responses^1–4^. However, microbiota enhance replication, transmission, or both with enteric viruses such as astroviruses, coxsackievirus B3, mouse mammary tumor virus, human and murine noroviruses, poliovirus, and reovirus strain T3SA+^5–12^. Binding to bacteria or bacterial surface polysaccharides directly enhance enteric viral infection by increasing cell attachment, virion stabilization, and viral co-infection^8,11,13–16^. Additionally, bacteria can modulate the host mucosal immune response to aid enteric viral infection^5,7,9^.

In most cases, microbiota inhibit viral infection outside of the intestinal tract but promote enteric viral infection. However, there are exceptions. For example, microbiota inhibit replication of murine norovirus (MNV) in the proximal small intestine by inducing local IFNλ responses, while simultaneously facilitating MNV replication in the distal intestine^17^. Also, microbiota depletion enhances rotavirus infection through reduced IL-22 expression, and segmented filamentous bacteria protect mice from rotavirus infection through effects on enterocyte migration^18–20^. Conversely, for non-enteric viruses, microbiota promote infection and virulence of Epstein-Barr virus and human immunodeficiency virus in humanized mice^21^. Therefore, microbiota can enhance infection with certain non-enteric viruses and inhibit infection with certain enteric viruses. Understanding mechanisms by which microbiota modulate viral infection is essential to fully comprehend the initiation of infection in the intestine.

Reovirus is a nonenveloped enteric virus with a genome of ten double-stranded RNA segments. Reovirus encodes three nonstructural proteins and eight structural proteins, including attachment protein σ1, an elongated fiber that binds distinct glycans followed by attachment to proteinaceous entry receptors^22–24^. The σ1 proteins from some reovirus strains bind glycans terminating in sialic acid (SA), which influences cell attachment, tropism, and pathogenesis^25,26^. Reovirus strains T3SA− and T3SA+ are isogenic viruses that differ by a single leucine-to-proline polymorphism at residue 204 in σ1 that confers SA binding to T3SA+. Relative to T3SA−, T3SA+ binds the cell surface more efficiently, infects cells that have low protein receptor expression^27^, and is more virulent in mice^28^. Based on the increased replication of T3SA+ in culture and *in vivo*, we used this strain over a decade ago for studies of the microbiota in perorally inoculated mice, in which we found that microbiota depletion inhibited T3SA+ replication and pathogenesis^11^.

Depending on the virus strain, after peroral inoculation of mice, reovirus can replicate in epithelial cells, M cells, or immune and other cell types within the lamina propria underlying the epithelium. However, intestinal sites of replication have been challenging to identify due to the limit of detection with viral antigen staining or electron microscopy^29–35^. Reovirus is unlikely to infect the epithelial cell layer from the apical side; instead, reovirus relies on M cells to gain access to the lamina propria followed by either replication in lamina propria cells or infection of intestinal epithelial cells from the basolateral side^31,32,35,36^.

Infection with reovirus and other viruses induces IFN responses that limit viral replication and disease^9,17,37–46^. Viral and bacterial molecules can be sensed by several mechanisms that culminate in production of type I (α/β), II (γ), or III (λ) IFNs. These cytokines bind distinct receptors (IFNAR, IFNGR, and IFNLR, respectively) to initiate signaling and expression of IFN-stimulated genes that have antiviral effector functions. The intestine has a compartmentalized innate response, with IFNs mediating different functions for different cell types^37,38,46^. While essentially all cells express IFNAR, IFNLR is predominantly expressed by mucosal epithelial cells that are preferentially protected from viral infection by IFNλ^37,39,42–44,47^. IFNα/β produced by intestinal immune cells restricts systemic dissemination of enteric viruses such as echovirus, MNV, and reovirus^9,38,40,41,45^. Reovirus strain T3D replicates inefficiently in lamina propria cells of immune-competent mice but replicates more efficiently in lamina propria cells and in extraintestinal tissues of *Ifnar^−/−^* mice that lack expression of the IFNα/β receptor^38,48^. However, in *Ifnlr1^−/−^* mice that lack expression of the IFNλ receptor, T3D tropism switches from lamina propria cells to epithelial cells that support robust replication, highlighting the importance of IFNλ restriction of viral infection in epithelial cells^38^.

In this study, we investigated mechanisms of microbiota enhancement or inhibition of enteric virus infection using a pair of isogenic reovirus strains. Microbiota depletion decreased replication of strain T3SA+ while paradoxically enhancing replication of strain T3SA−, which differs by one amino acid in the σ1 viral attachment protein. Both strains of reovirus bind to bacteria, which may facilitate delivery to target cells in the intestine. However, T3SA+ infected endothelial cells using a process dependent on SA and microbiota, whereas T3SA− infected intestinal epithelial cells where its replication was inhibited by microbiota-driven IFNλ responses. Therefore, a single amino acid polymorphism controls viral tropism and susceptibility to microbiota-driven host antiviral responses. These studies pinpoint viral glycan binding as a mediator of the interplay between enteric viruses and the host microbiota.

## RESULTS

### Reovirus strains are differentially affected by microbiota in the intestine

Microbiota depletion using antibiotics reduces replication and virulence of reovirus T3SA+ in perorally inoculated immunocompromised *Ifnar^−/−^*mice^11^. Therefore, similar to other enteric viruses (mouse mammary tumor virus, norovirus, and poliovirus)^5,7,8,11^, reovirus T3SA+ benefits from microbiota that occupy its infection niche. However, we found that replication of other reovirus strains was increased in microbiota-depleted mice. Conventional or antibiotic-treated C57BL/6 mice were perorally inoculated with 10^8^ PFU of reovirus strain T1L, followed by assessment of viral titers at 4 days post-infection (dpi) by plaque assay. Microbiota-depleted mice had 15-fold higher T1L fecal shedding (Figure 1A). We observed similar results for T1L infection of microbiologically sterile germ-free C57BL/6 mice. Germ-free mice that were colonized with fecal bacteria (GF-Conv) prior to infection with T1L had a 50-fold decrease in viral shedding, further demonstrating an inhibition of viral replication in the presence of microbiota (Figure 1A). Because microbiota can enhance or inhibit acute MNV-1 infection in different regions of the intestine^17^, we assessed T1L titers across the small intestine of perorally inoculated conventional and germ-free mice. We found that T1L titers in germ-free mice were increased in all regions of the intestine, suggesting that bacteria inhibit T1L infection broadly (Figure 1B). Next, we investigated whether the opposing effects of the microbiota on reovirus T3SA+ in *Ifnar^−/−^* mice and T1L in immune-competent mice were due to differences in the viral strains or the immune status of the mice. We found that titers of T3SA+ following peroral inoculation of conventional, germ-free, or antibiotic-treated immune-competent C57BL/6 mice were reduced in mice with depleted microbiota (Figure 1C, left). These results suggest that differences in the viral strains dictate microbiota reliance vs. inhibition.

**Figure 1.**
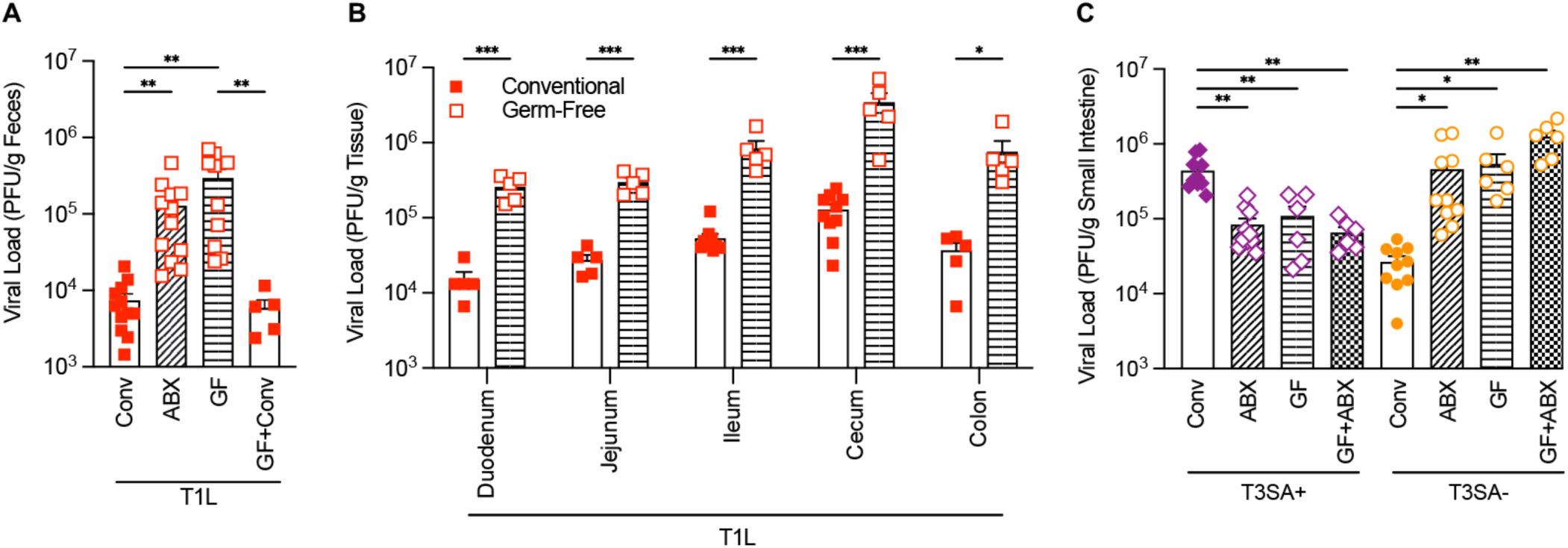
Microbiota depletion differentially affects replication of distinct reovirus strains in the intestine. (A) Microbiota depletion increases reovirus T1L shedding. Adult conventional (Conv) C57BL/6 mice were treated with or without antibiotics (ABX) and germ-free (GF) C57BL/6 mice were treated with or without feces from conventional mice to establish microbiota (GF+Conv) prior to peroral inoculation with 10^8^ PFU of T1L. Viral titers in feces at 4 dpi were determined by plaque assay using L929 cells (n=5-12 mice per condition).(B) Microbiota inhibit reovirus T1L infection throughout the intestine. Conventional or germ-free C57BL/6 mice were perorally inoculated with 10^8^ PFU of T1L. Viral titers in small intestine at 4 dpi were determined by plaque assay (n=5-10 mice per condition). (C) Microbiota promote infection by T3SA+ but inhibit infection by T3SA−. Adult conventional (Conv) C57BL/6 mice were treated with or without antibiotics (ABX) and germ-free (GF) C57BL/6 mice were treated with or without antibiotics (GF+ABX) prior to peroral inoculation with 10^8^ PFU of T3SA+ or T3SA−. Viral titers in small intestine at 4 dpi were determined by plaque assay (n=6-10 mice per condition). Columns and error bars show mean ± SEM from at least two independent experiments. Filled symbols indicate microbiota-replete mice and open symbols and striped columns indicate microbiota-depleted mice. Significance was assessed using multiple unpaired t tests. *, *P <* 0.05; **, *P <* 0.01; ***, *P <* 0.001.

Since reovirus strain differences influence the effects of microbiota on viral infection for strains T1L vs. T3SA+, we sought to identify the relevant viral determinants. T3SA+ has nine genome segments derived from T1L and a σ1-encoding gene segment from a distinct strain, T3C44-MA^27^, implicating differences in the σ1 protein in differential microbiota effects. The T1L and T3SA+ σ1 proteins have 21% amino acid sequence identity^49^, which complicates efforts to identify the relevant mutations. However, T3SA+ is derived from isogenic strain T3SA−, which differs from T3SA+ by a single leucine-proline polymorphism at residue 204 in the σ1 protein that modulates SA binding^27^. Pro204 in σ1, which confers SA binding to T3SA+, was selected during serial passage of T3SA− in murine erythroleukemia (MEL) cells that have low protein receptor expression with efficient infection reliant on SA-binding capacity^27,50^. To determine whether the P204L polymorphism in σ1 dictates microbiota reliance vs. inhibition, we compared titers of T3SA+ and T3SA− in perorally inoculated conventional, antibiotic-treated, germ-free, or antibiotic-treated germ-free C57BL/6 mice. We found that T3SA+ titers were reduced in all mice with reduced or absent microbiota. However, T3SA− titers were increased in all mice with reduced or absent microbiota (Figure 1C). Therefore, a single amino acid that dictates SA-binding capacity controls susceptibility to microbiota for these reovirus strains. Additionally, titers for a given virus were equivalent in germ-free mice and germ-free mice treated with antibiotics, suggesting that the effects are attributable to the intestinal microbiota and not off-target effects of antibiotic treatment (Figure 1C).

### T3SA+ and T3SA− reovirus strains bind to bacteria

Bacteria produce SA and SA-like molecules in addition to many other glycans. Therefore, it is possible that T3SA+ and T3SA− differ in binding to bacteria, which may influence microbiota reliance vs. inhibition. Both type 1 (T1L) and type 3 (T3D) reovirus strains bind to bacteria^16^, suggesting that bacterial binding is not dependent solely on SA and may involve reovirus capsid proteins other than σ1. For other enteric viruses, including noroviruses and picornaviruses, viral binding to bacteria using bacterial-surface glycans can have a range of effects, including increased attachment to host cells, increased virion stability, and increased delivery of multiple virions to cells^11,14–16,51^. Since T3SA+ and T3SA− have different SA-binding capacities ^52,53^, and binding to bacteria may aid viral delivery to host cells, we quantified the relative bacterial binding efficiencies of these strains. First, we used confocal microscopy of fluorescently-labeled T3SA+ or T3SA− virions incubated with DAPI-stained Gram-negative bacteria (*E. coli*) or Gram-positive bacteria (*B. cereus)*. We observed both T3SA+ and T3SA− virions (green) bound to *E. coli* and *B. cereus* (blue) (Figure 2A). Next, we quantified viral binding to bacteria using a flow cytometry assay with unlabeled T3SA+ or T3SA− virions to ensure that fluorescent labeling of virions would not affect potential binding sites. Both reovirus strains were capable of binding to *E. coli* and *B. cereus*, although T3SA+ bound to an increased percentage of both bacterial strains relative to T3SA− (Figure 2B). Thus, T3SA+ and T3SA− reovirus strains bind to bacteria, which may influence their delivery to host cells and infection in the intestine.

**Figure 2.**
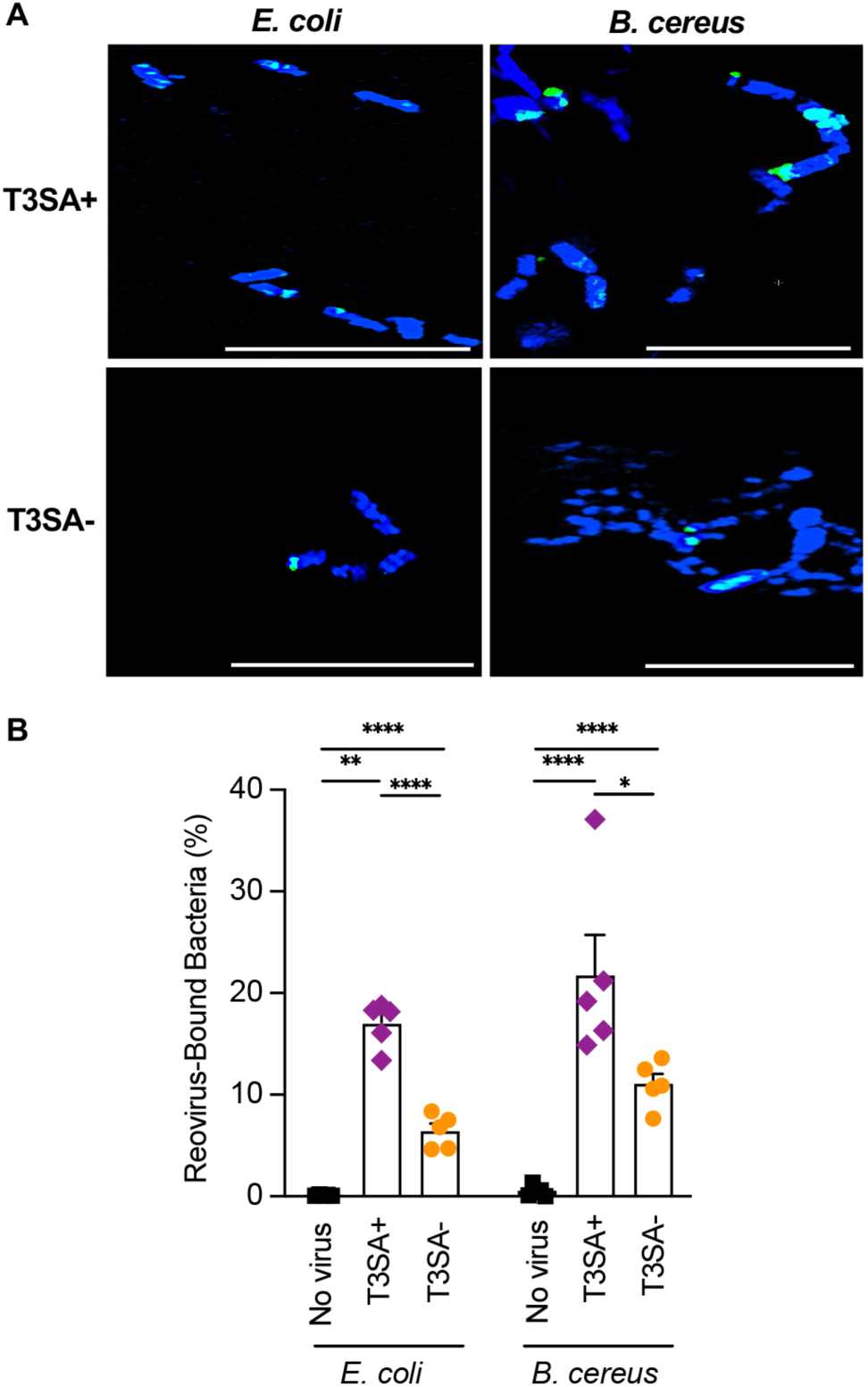
Reovirus strains T3SA+ and T3SA− bind to bacteria. (A) *E. coli* or *B. cereus* were incubated with Fluorescently-labeled T3SA+ or T3SA− at room temperature for 15 minutes. Bacteria were stained with DAPI and imaged using confocal microscopy. Scale bar, 10 μm. (B) Quantification of reovirus-bound bacteria using flow cytometry. *E. coli* or *B. cereus* were incubated with buffer (No virus), T3SA+, or T3SA− at 10^3^ viral particles/bacterial cell at room temperature for 30 min. Bacteria were stained with reovirus-specific polyclonal antiserum and anti-rabbit Alexa-488. Reovirus bound-bacteria were quantified using flow cytometry. The percentage of bacteria bound to 488-labeled reovirus was determined from 10^5^ total events. The data are presented as the mean ± SEM from 5 biological replicates in two independent experiments (each biological replicate is the average of sample analyzed in triplicate). Significance was assessed using multiple unpaired t tests. *, *P <* 0.05; **, *P <* 0.001; ****, *P <* 0.0001.

### Microbiota enhancement of T3SA+ infection is SA-dependent

The reovirus σ1 protein determines the viral serotype and specific glycans used for initial cell attachment. The glycan-binding region of serotype 3 σ1 protein is centered on residue 204, which is located in the body domain of the protein (Figure 3A)^25,27,53,54^. To determine whether microbiota effects on reovirus strains are due to SA binding or are due to consequences of the residue 204 polymorphism independent of SA binding, we engineered alternative mutations to ablate SA binding by targeting distinct amino acids, N198 and R202 (Figure 3B). First, we compared SA-binding capacity of wild-type and mutant reovirus strains using hemagglutination (HA) assays. HA titers were significantly lower for T3SA− and the additional non-SA-binding mutants, T3SA− (N198D) and T3SA− (R202W), compared with T3SA+ or prototype reovirus strain T3D (Figure 3C). Next, we tested whether the SA-binding mutant viruses have replication defects following infection of L929 cells, which are highly susceptible to all reovirus types, and murine erythroleukemia (MEL) cells, which are only susceptible to infection with SA-binding reovirus strains^50^. In L929 cells, all reovirus strains replicated efficiently and produced comparable viral titers at all time points tested. However, the reovirus strains incapable of binding SA did not replicate efficiently in MEL cells and produced viral titers at 24 hpi that were more than 10-fold lower than SA-binding strains T3SA+ and T3D (Supplemental Figure 1). Finally, to examine contrasting effects of microbiota on SA-binding and non-SA-binding reovirus strains, we quantified titers of T3SA+, T3SA− (N198D), or T3SA− (R202W) in the small intestines of perorally inoculated conventional or antibiotic-treated mice. As expected, T3SA+ titers were reduced in antibiotic-treated mice, while titers of reovirus strains incapable of binding to SA, T3SA− (N198D) and T3SA− (R202W), were significantly increased, similar to results with T3SA− (compare Figure 3D and Figure 1C). These results suggest that the effect of the intestinal microbiota on reovirus infection is dependent on SA-binding capacity of the reovirus strain.

**Figure 3.**
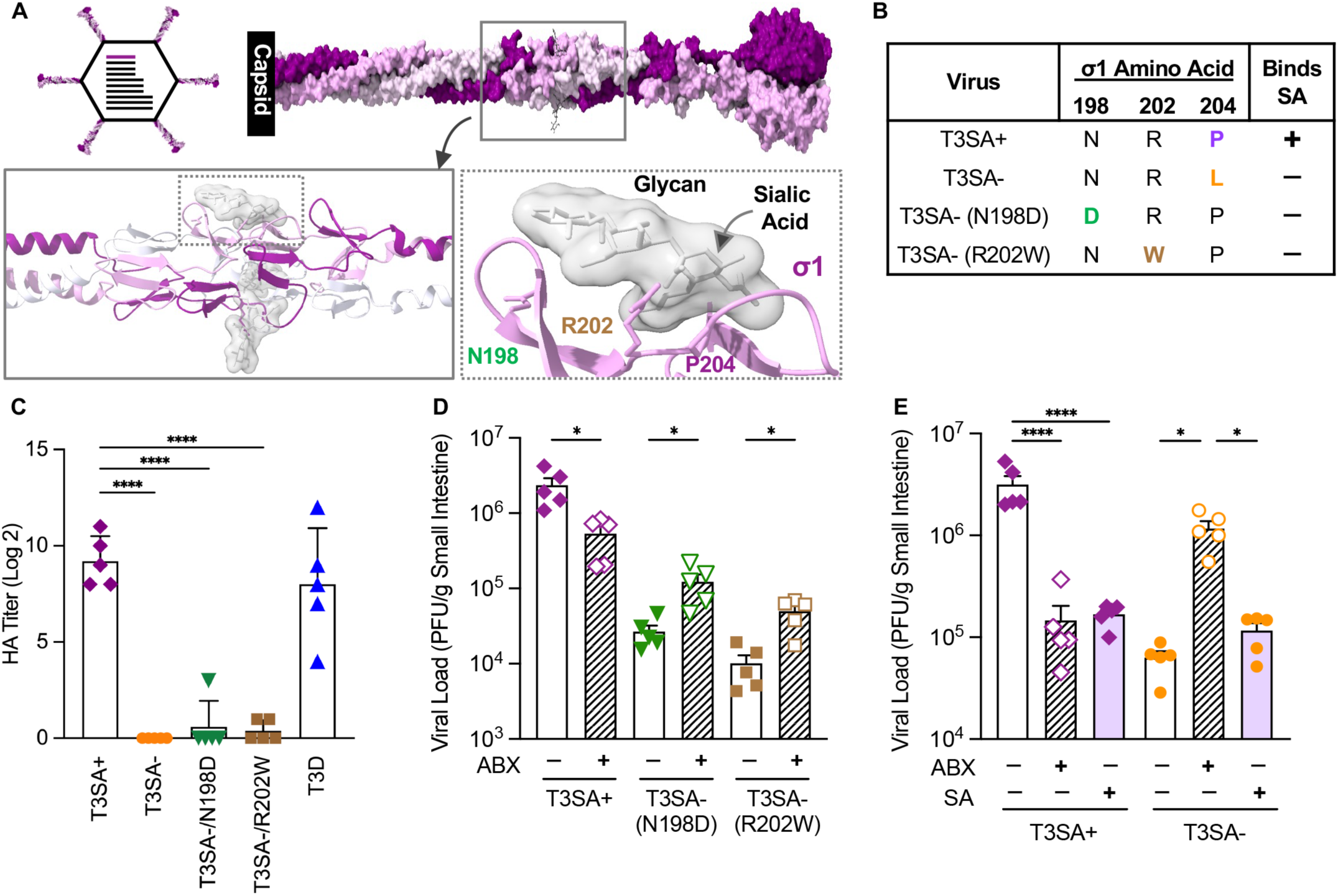
Sialic acid binding capacity of the reovirus σ1 attachment protein dictates microbiota reliance vs. susceptibility. (A) Top left: Schematic of reovirus T3SA+ particles. This strain contains nine genome segments (black lines) from T1L and the S1 (σ1 protein-encoding) genome segment (purple line) from a type 3 strain. The σ1 attachment protein (purple stalk) is shown at the icosahedral vertices of the virion. Top right: Full-length surface-shaded model of the trimeric T3SA+ σ1 protein, which binds to cell-surface glycans that terminate in SA using sequences in the central body domain (boxed). Bottom: Insets show enlarged σ1 regions near the SA-binding site of T3SA+, with relevant amino acids highlighted. Structure model of the trimeric σ1 protein [derived from PDB accession numbers 6GAP for amino acids 27–243^54^ and 3S6X for amino acids 244– 455^25,54^. (B) The σ1 amino acid changes (orange, green, or brown) that were engineered to ablate T3SA+ binding to SA. (C) SA-binding capacity of reovirus strains. Purified virions were mixed with erythrocytes, and hemagglutination (HA) titers were determined following incubation at 4°C for 3 h. Results are shown as mean ± SEM. Significant differences from T3SA+ were assessed using one-way ANOVA and Dunnett’s test. (D) Ablation of SA-binding by specific mutations in σ1 renders viruses susceptible to microbiota-mediated infection inhibition. Conventional C57BL/6 mice were treated with or without antibiotics (ABX) prior to peroral inoculation with 10^8^ PFU of T3SA+ or 5 × 10^7^ PFU of T3SA−/N198D or T3SA−/R202W. Viral titers in small intestine at 4 dpi were determined by plaque assay (n=5 mice per condition). (E) Exogenous SA inhibits T3SA+ infection to the same extent as microbiota depletion. Conventional C57BL/6 mice were treated with or without antibiotics (ABX) or with exogenous *N*-acetylneuraminic acid/sialic acid (SA). SA (1 mg) was adminstered orally twice/day for 2 d prior to inoculation as well as administered in the drinking water at a concentration of 1% throughout the experiment. Mice were perorally inoculated with 10^8^ PFU of T3SA+ or T3SA−, and viral titers in small intestine at 4 dpi were determined by plaque assay (n=5 mice per condition, two independent experiments). (D and E) Data are shown as mean ± SEM. Significant differences were assessed using ordinary one-way ANOVA with Fisher’s LSD test. *\*, P <* 0.05; **, *P <* 0.01; ***, *P <* 0.001; ****, *P <* 0.0001.

Microbiota influence SA levels in the intestine, which may affect viral infection^55–57^. Since intestinal bacteria use SA as a nutrient source, depletion of microbiota increases levels of free SA in the intestinal lumen by 20-70-fold^57^. We speculated that excess free SA in antibiotic-treated mice could bind to T3SA+ virions prior to cell attachment and inhibit infection. To address this possibility, we gavaged mice perorally with PBS, antibiotics, or orally administered exogenous SA prior to inoculation with T3SA+ or T3SA− and assessed viral titers in the small intestine at 4 dpi. Compared with PBS-treated mice, SA administration decreased T3SA+ titers by 20-fold, comparable to titers in antibiotic-treated mice (Figure 3E). In contrast, SA administration did not alter T3SA− titers in the intestine, indicating that exogenous SA specifically inhibited the SA-binding strain. These results suggest that bacteria-mediated enhancement of reovirus T3SA+ infection is dependent on viral SA binding and that excess SA in the intestine of microbiota-depleted mice contributes to reduced replication of T3SA+.

### Reovirus strains T3SA+ and T3SA− have distinct cellular tropism within the intestine

Reoviruses have broad cell and tissue tropism, which is influenced by viral strain and SA-binding capacity^53^. We found that microbiota-enhancement of T3SA+ infection of the small intestine also is dependent on SA binding, suggesting that microbiota influence replication efficiency or tropism in the intestine. Previous immunohistochemistry studies found few reovirus-infected cells, located primarily in the Peyer’s patches of the small intestine of adult wild-type mice, despite detectable tissue titers^38^. Therefore, we used a hybridization chain reaction RNA fluorescence *in situ* hybridization (HCR-FISH) assay, which has enhanced sensitivity relative to immunohistochemistry^58^. Conventional or antibiotic-treated C57BL/6 mice were mock-infected or perorally inoculated with T3SA+ or T3SA−. At 4 dpi, Swiss-rolled small intestinal tissues were processed for HCR-FISH using probes designed to detect reovirus viral RNA (vRNA). Following infection of conventional mice, there were numerous T3SA+ vRNA-positive cells localized within the lamina propria of intestinal villi (Figure 4A). Antibiotic-treated mice had fewer T3SA+ vRNA-positive cells overall, but infected cells remained localized to the lamina propria. In contrast, T3SA− vRNA-positive cells were rare but almost exclusively found in intestinal epithelial cells of both conventional and antibiotic-treated mice. We quantified the number of vRNA-positive villi from representative confocal micrographs obtained throughout the small intestine. In concordance with viral titers in the small intestine (Figures 1C and 3E), the percentage of infected villi in conventional mice inoculated with T3SA+ was significantly higher than in antibiotic-treated mice, whereas the findings were inversed for mice inoculated with T3SA− (Figure 4B). Next, we quantified reovirus-positive foci in epithelial cells or lamina propria cells of intestinal villi relative to the average number of positive foci in 100 representative villi of immune-competent, conventional mice. We found that T3SA+ and T3SA− had strikingly different cell tropism. Following inoculation of conventional or antibiotic-treated mice, T3SA+ predominantly infected cells in the lamina propria, whereas T3SA− infected cells primarily in the epithelial layer (Supplemental Table 1, WT mice). These data show that reovirus strains T3SA+ and T3SA− have distinct intestinal cell tropism and that the tropism of each virus was not altered by microbiota depletion.

**Figure 4.**
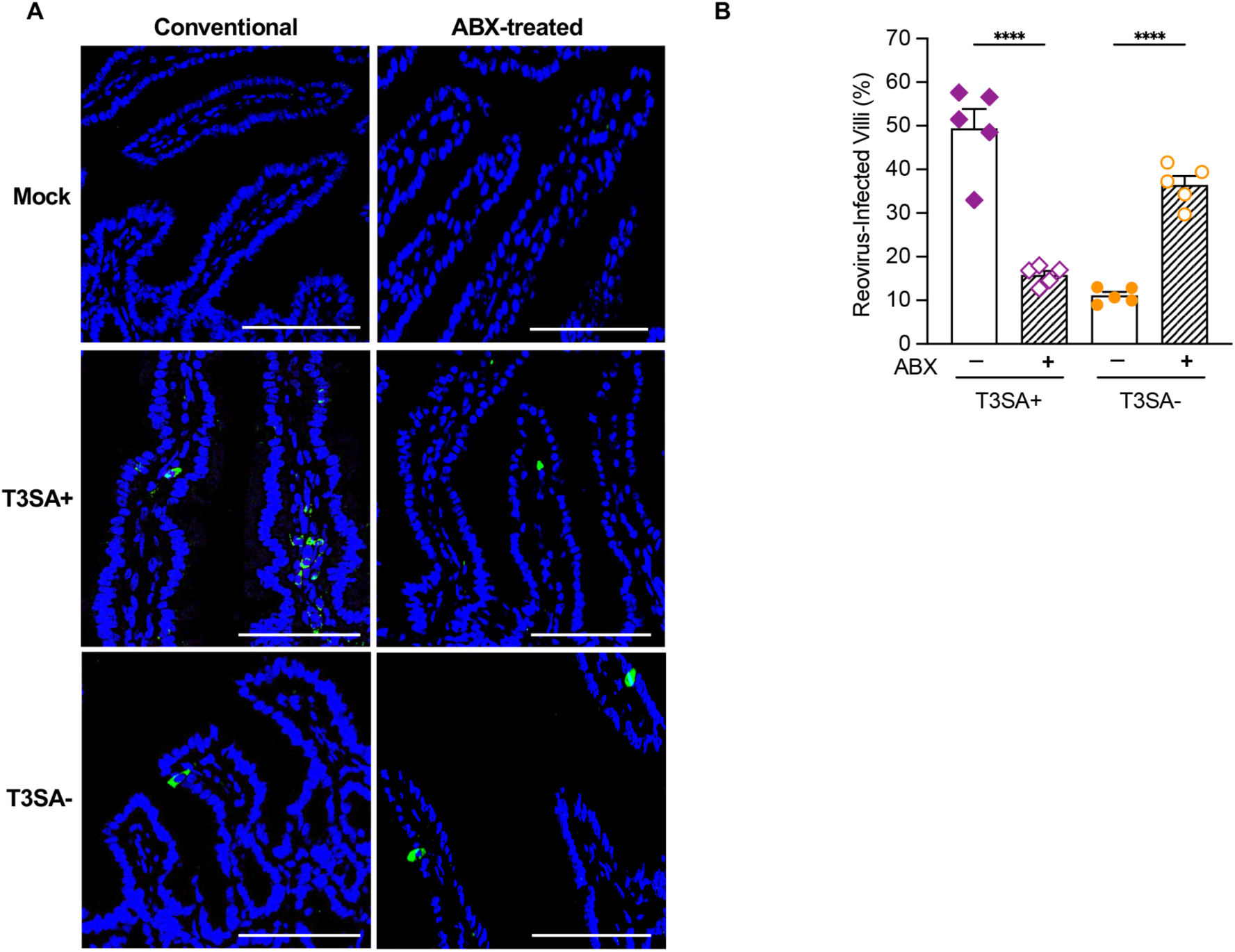
Reovirus strains T3SA+ and T3SA− have distinct cell tropism and infection efficiency in the intestine. Conventional C57BL/6 mice were treated with or without antibiotics (ABX) prior to peroral inoculation with PBS (Mock) or 10^8^ PFU of T3SA+ or T3SA−. At 4 dpi, the small intestine was harvested and Swiss rolled for tissue sectioning and staining with an HCR-FISH assay using probes to detect S4 viral RNA (vRNA)/infected cells (green) and DAPI to detect nuclei (blue). (A) Representative images show substantial T3SA+ infection of cells within the lamina propria, which is reduced in ABX-treated mice and sporadic T3SA− infection of intestinal epithelial cells, which is increased in ABX-treated mice. Scale bars, 100 μm. (B) Quantification of reovirus-infected villi from confocal images of 100 villi per mouse. Data are shown as mean ± SEM (n=5 mice). Significance was assessed using ordinary one-way ANOVA with Fisher’s LSD test. ****, *P <* 0.0001.

### Reovirus T3SA+ infects intestinal endothelial cells in a SA-dependent manner

Following peroral inoculation, type 1 reovirus strains primarily infect intestinal epithelial cells, while SA-binding type 3 strains preferentially infect cells within the lamina propria of intestinal villi (Figure 4A)^38,59^. T3SA+ also infects polarized human brain microvascular endothelial cells *in vitro*^36^. To identify reovirus-infected cells in the intestine, we perorally inoculated C57BL/6 mice with T3SA+ or T3SA− and collected intestinal tissue sections at 4 dpi and conducted HCR immunofluorescence (IF) and RNA-FISH analyses for simultaneous high-resolution protein and vRNA imaging *in vivo*. Tissues were stained with antibodies specific for mouse immune cells (CD45) and endothelial cells (CD31) and corresponding secondary fluorescent probes followed by RNA-FISH probes to detect reovirus vRNA. We did not observe colocalization of T3SA+ vRNA with CD45+ immune cells (Figure 5A). However, T3SA+ vRNA colocalized with CD31+ endothelial cells, while T3SA− vRNA did not (Figure 5B). These results suggest that reovirus SA-binding capacity facilitates infection of intestinal endothelial cells. To test this hypothesis, we used primary small intestine endothelial cells isolated from immune-competent mice. Primary endothelial cells or control highly susceptible mouse L929 cells were inoculated with T3SA+ or T3SA−, and viral infectivity was quantified at 18 hpi. Both T3SA+ and T3SA− efficiently infected L929 cells (Figure 5B). Infection of endothelial cells was less efficient than infection of L929 cells, but T3SA+ produced significantly higher yields in endothelial cells than those produced by T3SA− (Figure 5B). To examine whether T3SA+ SA-binding capacity influenced endothelial cell infection, we pre-treated endothelial cells with neuraminidase to remove cell-surface SA or pre-incubated virus stocks with excess SA to saturate binding sites prior to endothelial cell infection and viral titer analysis. Treatment of cells with neuraminidase or virus with excess SA significantly decreased infection of endothelial cells by T3SA+ but had no effect on infection of these cells by T3SA− (Figure 5C). Collectively, these data show that reovirus SA-binding capacity facilitates infection of intestinal endothelial cells.

**Figure 5.**
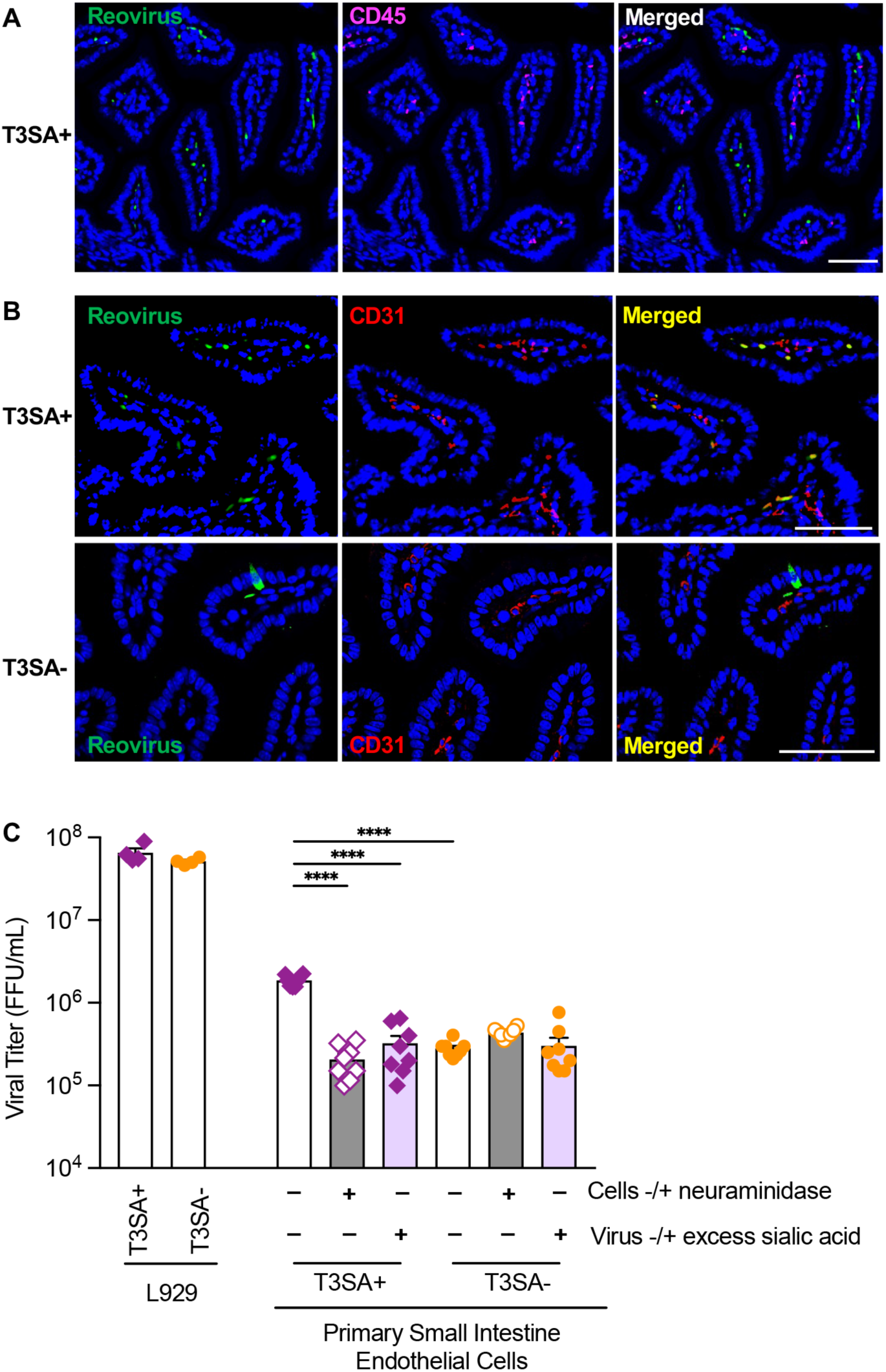
Reovirus T3SA+ infects intestinal endothelial cells. (A) T3SA+ vRNA does not colocalize with CD45, which marks immune cells. C57BL/6 mice were perorally inoculated with 10^8^ PFU of T3SA+. At 4 dpi, the small intestine was harvested and processed for simultaneous protein and RNA detection using HCR IF and RNA-FISH, respectively. Shown are representative images of intestinal villi using probes specific for reovirus (green) and CD45 (magenta). DAPI-stained nuclei are shown in blue. Scale bar, 100 μm. (B) T3SA+ but not T3SA− vRNA colocalizes with CD31, which marks endothelial cells, in the lamina propria. The experiment was conducted as in panel A, but both T3SA+ and T3SA− were used for independent infections and tissues were stained with CD31 antibody. Shown are representative images of intestinal villi using probes specific for reovirus (green) and CD31 (red). Scale bars, 100 μm. (C) T3SA+ infects primary mouse intestinal endothelial cells more efficiently than does T3SA− in an SA-dependent manner. Highly susceptible L929 cells were infected with T3SA+ or T3SA− at an MOI of 0.01. Infected cells were quantified at 18 hpi by FFU assay. Primary small intestine endothelial cells were treated with or without 80 mU/ml *Clostridium perfringens* neuraminidase to remove cell-surface SA prior to infection with T3SA+ or T3SA− stocks that were pre-incubated with or without SA (10 μg/mL) at an MOI of 0.01. Infected cells were quantified at 18 hpi by FFU assay. The data are presented as the mean FFU/mL ± SEM from two independent experiments with data points representing individual samples (n=4 for L929 cells and n=8 for endothelial cells). Significance was assessed using Brown-Forsythe and Welch ANOVA tests. ****, *P <* 0.0001.

### Lack of SA binding restricts T3SA− infection to intestinal epithelial cells where replication is inhibited by microbiota-driven host IFNλ responses

Type-III IFN (IFNλ) responses control intestinal infection of several enteric viruses, particularly those that replicate in epithelial cells, since these cells express the INFλ receptor, IFNLR1^17,37,41–44,47,60^. Reovirus strains T1L and T3D shed in the stool more efficiently and produce higher viral titers in the intestine of *Ifnlr1^−/−^* mice^38,44^. Because IFNλ responses inhibit viral replication in epithelial cells^37,42–44,47^, and microbiota can modulate IFN responses^43,61–64^, we tested whether T3SA+ and T3SA− are sensitive to IFNλ responses by infecting microbiota-replete or microbiota-depleted *Ifnlr1^−/−^* mice. Conventional or antibiotic-treated immune-competent C57BL/6 or *Ifnlr1^−/−^*mice were perorally inoculated with T3SA+ or T3SA−. At 4 dpi, the entire small intestine was harvested for analysis of viral titer or Swiss-rolled and processed for HCR RNA-FISH to detect vRNA-containing cells. We found that the lack of IFNλ sensing in *Ifnlr1^−/−^* mice had no effect on infection by T3SA+. Titers of this strain in *Ifnlr1^−/−^* mice were equivalent to those in immune-competent mice (Figure 6A), microbiota depletion inhibited replication (Figures 6A, 6B, and 6C), and replication occurred primarily in lamina propria cells (Figures 6B and 6D and Supplemental Table 1). In contrast, lack of IFNλ sensing in *Ifnlr1^−/−^* mice increased T3SA− titers by 12-fold and increased the number of vRNA-containing cells (Figures 6A, 6B, and 6C). In contrast to findings with T3SA+, the percentage of villi infected with T3SA− was higher in *Ifnlr1^−/−^*mice relative to immune-competent mice (compare Figures 4B and 6C and Supplemental Table 1). While replication of T3SA− remained restricted primarily to epithelial cells in *Ifnlr1^−/−^* mice (Figures 6B, 6D, Supplemental Table 1), T3SA− infection was increased in microbiota-replete mice to that in microbiota-depleted mice. For T3SA−, lack of IFNλ sensing phenocopied microbiota depletion in immune-competent animals, and microbiota depletion in *Ifnlr1^−/−^*mice did not increase infection (Figure 6). These findings suggest that microbiota prime the intestinal epithelium for IFNλ production, which inhibits T3SA− replication in epithelial cells. Conversely, T3SA+ is unaffected by microbiota-driven IFNλ restriction due to the preferential infection of IFNλ-resistant endothelial cells by this strain. Overall, these results suggest that SA-binding capacity determines viral tropism, which influences microbiota sensitivity by the action of cell-specific innate immune responses.

**Figure 6.**
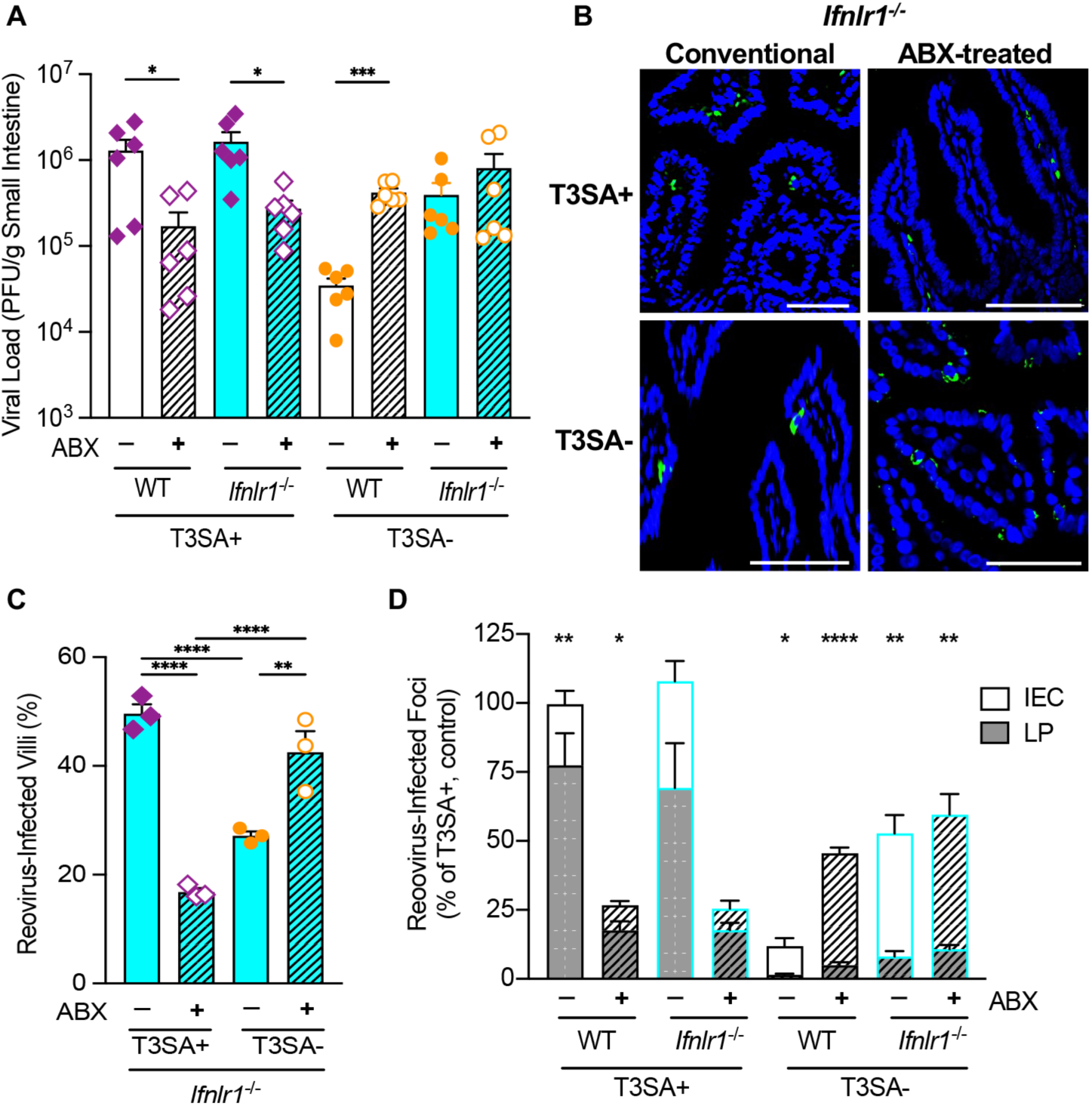
Reovirus T3SA− infection of intestinal epithelial cells is restricted by microbiota-induced IFNλ responses. (A) Microbiota depletion or ablation of IFNλ response increases T3SA− infection. Conventional C57BL/6 WT or *Ifnlr1^−/−^* mice were treated with or without antibiotics (ABX) prior to peroral inoculation with 10^8^ PFU of T3SA+ or T3SA−. At 4 dpi, the small intestine was harvested for determination of viral titers by plaque assay. The data are presented as mean ± SEM (n=6 mice per condition, two independent experiments). (B) Representative images of cells infected with T3SA+ and T3SA− in the intestines of *Ifnlr1^−/−^* mice. Experiments were conducted as in panel A except that tissues were processed for HCR-FISH detection of vRNA (green) using confocal microscopy. DAPI-stained nuclei are shown in blue. Scale bars, 100 μm. (C) Quantification of T3SA+ and T3SA− infection in intestines of *Ifnlr1^−/−^* mice. The percentages of reovirus-infected villi were determined from images of 100 villi per mouse using vRNA HCR. The data are presented as mean ± SEM (n=3 mice). (D) Quantification of reovirus-infected foci in intestinal epithelial cells (IEC) or lamina propria cells (LP). Micrograph images were enumerated for vRNA-stained cells using Fiji from 100 villi per mouse. The data are presented as percentage of total (IEC+LP) reovirus-positive cells in each mouse relative to T3SA+ infection of conventional WT mice. (A,C-D) Significance was assessed using (A) Brown-Forsythe and Welch ANOVA tests, (C) ordinary one-way ANOVA with Fisher’s LSD test, (D) multiple unpaired t tests.*, *P <* 0.05; **, *P <* 0.001; ****, *P <* 0.0001.

## DISCUSSION

Microbiota influence viral infection by diverse mechanisms. In general, enteric viruses benefit from intestinal microbiota that occupy their infection site, while respiratory or systemic viruses that infect extraintestinal tissues are inhibited by microbiota due to stimulation of host antiviral responses^3,4,12,21,41,43,65–67^. Fitting with these themes, we found that microbiota enhance infection by reovirus strain T3SA+ (Figures 1, 3, 4, 6)^11^. However, this strain is an outlier among *Reovirales* viruses, as reovirus strains T1L and T3SA− produce lower viral titers in the presence of microbiota (Figure 1), and intestinal bacteria inhibit rotavirus infection^18–20^. Reovirus strains T3SA+ and T3SA− differ by a single amino acid in the σ1 attachment protein that regulates SA binding. Using these otherwise isogenic reovirus strains, we found that SA binding facilitates reovirus infection of intestinal endothelial cells rather than epithelial cells, which allows T3SA+ to avoid microbiota-driven IFNλ responses in epithelial cells that limit T3SA− infection. Therefore, viral tropism dictates how the intestinal microbiota influence enteric viral infection.

Replication of enteric viruses can be enhanced by virus binding to bacteria and bacterial surface glycans, which can facilitate viral attachment to host cells, delivery of multiple virions to the cytoplasm, and stabilization of virions to avoid premature disassembly^8,11,14,15,68,69^. For example, poliovirus binds bacterial lipopolysaccharide, which stabilizes virions and aids viral receptor binding^11,14^, and noroviruses bind histo-blood group antigens expressed on certain bacteria, which likewise enhances viral stability and attachment to host cells^8,68,69^. Here, we found that both T3SA+ and T3SA− bind to bacteria (Figure 2), which aligns with prior data showing that reovirus strains T1L and T3D bind to and are stabilized by bacteria^16^. Reovirus bound to bacteria was readily transferred to host cells, suggesting that low-affinity interactions facilitate viral delivery to host cells^16^. We found that T3SA+ bound to bacteria more efficiently than T3SA− (Figure 2B), suggesting that SA-binding capacity influences bacteria binding. Although strain-specific differences in reovirus-bacteria interactions are the focus of our current research, the role of bacteria binding during reovirus infection in the intestine is unclear. Nonetheless, it is possible that bacteria facilitate delivery of reovirus to host cells within the intestine, which may explain how the microbiota enhance T3SA+ infection.

In addition to possible bacteria-mediated delivery of reovirus virions to host cells in the intestine, T3SA+ also likely benefits from microbiota-mediated sequestration of free SA in the intestinal lumen to aid SA-dependent viral attachment to host cells. SAs derived from host cells or mucins can be used by the microbiota as a source of carbon and nitrogen^55,56^. SA levels in the intestinal lumen are low in conventional mice but elevated in microbiota-depleted or germ-free mice due to reduced consumption by the microbiota^57^. These elevated levels of SA in microbiota-depleted mice can be recapitulated by oral administration of exogenous SA to microbiota-replete mice^57^. We found that T3SA+ titers were reduced in microbiota-replete mice treated with exogenous SA, phenocopying microbiota depletion, while T3SA− titers were unchanged by SA treatment (Figure 3E). These results suggest that, in addition to aiding in the delivery of virions to host cells, microbiota facilitate infection of T3SA+ by maintaining low SA concentrations in the lumen, thus limiting premature occupation of viral SA-binding sites that facilitate host cell attachment.

Enteric viruses infect different types of cells within the intestine. Viral infection and tropism is influenced by intestinal microbiota by direct effects on cell attachment and indirect effects on immune responses^9,17,38,39,65,70^. For reoviruses, the efficiency of intestinal infection depends on host age, immune status, and viral strain. Type 1 reovirus strains generally infect intestinal epithelial cells^59^, whereas type 3 strains generally infect lamina propria cells and Peyer’s patches^38,71,72^. SA binding enhances reovirus infection efficiency in mice following peroral or intracranial inoculation^28,53,73^. However, the cell tropism of T3SA+ and T3SA− in the mouse intestine was largely unknown. Using a sensitive HCR RNA-FISH assay to detect reovirus RNA, we found a striking difference in the tropism of these strains. T3SA+ primarily infected cells within the lamina propria, while T3SA− primarily infected epithelial cells. Similar to viral titers in intestinal tissues, the number of infected foci was altered by microbiota depletion, but cell tropism did not change. Additionally, we found that T3SA+ colocalized with CD31+ endothelial cells and, concordantly, infection of primary intestinal endothelial cells was dependent on viral SA-binding capacity. These findings demonstrate that reovirus cell tropism in the intestine is strongly influenced by SA binding.

Since T3SA− replicates in intestinal epithelial cells, we investigated whether epithelial cell-specific innate immune responses stimulated by the microbiota underlie the increased infection of T3SA− in microbiota-depleted mice. IFNλ responses inhibit replication of viruses at epithelial surfaces^37,42–44,47^, and microbiota can stimulate IFN responses^43,61,63,64,74^. IFNλ responses inhibit replication of reovirus T1L^44^ and T3D in epithelial cells and influence T3D tropism. T3D primarily infects lamina propria cells in wild-type or *Ifnar^−/−^* mice, but in *Ifnlr1^−/−^* mice, T3D infects epithelial cells^38^. We found increased T3SA− infection in epithelial cells of *Ifnlr1^−/−^* mice, mimicking effects of microbiota depletion. Microbiota depletion did not increase T3SA− replication in *Ifnlr1^−/−^*mice, suggesting that the reduced replication of T3SA− in epithelial cells is attributable to microbiota-driven IFNλ responses. T3SA+ replication was not altered in *Ifnlr1^−/−^* mice, suggesting that T3SA+ is resistant to the antiviral effects of IFNλ *in vivo*, likely due to its non-epithelial replication site in endothelial cells. Collectively, these data suggest that T3SA− is subjected to microbiota-driven innate immune responses in epithelial cells, which are alleviated by microbiota depletion.

Coupled with prior studies, our data support a unified model (Figure 7) in which microbiota promote or inhibit enteric virus infection at least in part based on viral tropism within the intestine. We hypothesize that, in the intestinal lumen, reovirus strains bind bacteria with varying affinity, which facilitates delivery of virions to the lamina propria. Subsequently, viral SA-binding capacity dictates reovirus tropism. SA-binding strains such as T3SA+ infect cells in the lamina propria that are resistant to IFNλ responses, while non-SA-binding strains such as T3SA− infect intestinal epithelial cells that are sensitive to microbiota-driven IFNλ responses. This model provides an explanation for the observation that several viruses in the *Reovirales* are outliers in microbiota effects on enteric viruses. Reovirus strains T1L and T3SA− as well as certain rotavirus strains infect intestinal epithelial cells and are inhibited by microbiota-driven host antiviral responses, whereas many other enteric viruses, including reovirus T3SA+, infect lamina propria cells and are not subjected to these antiviral responses. These results provide a mechanistic explanation for disparate effects of microbiota depletion on enteric viruses.

**Figure 7.**
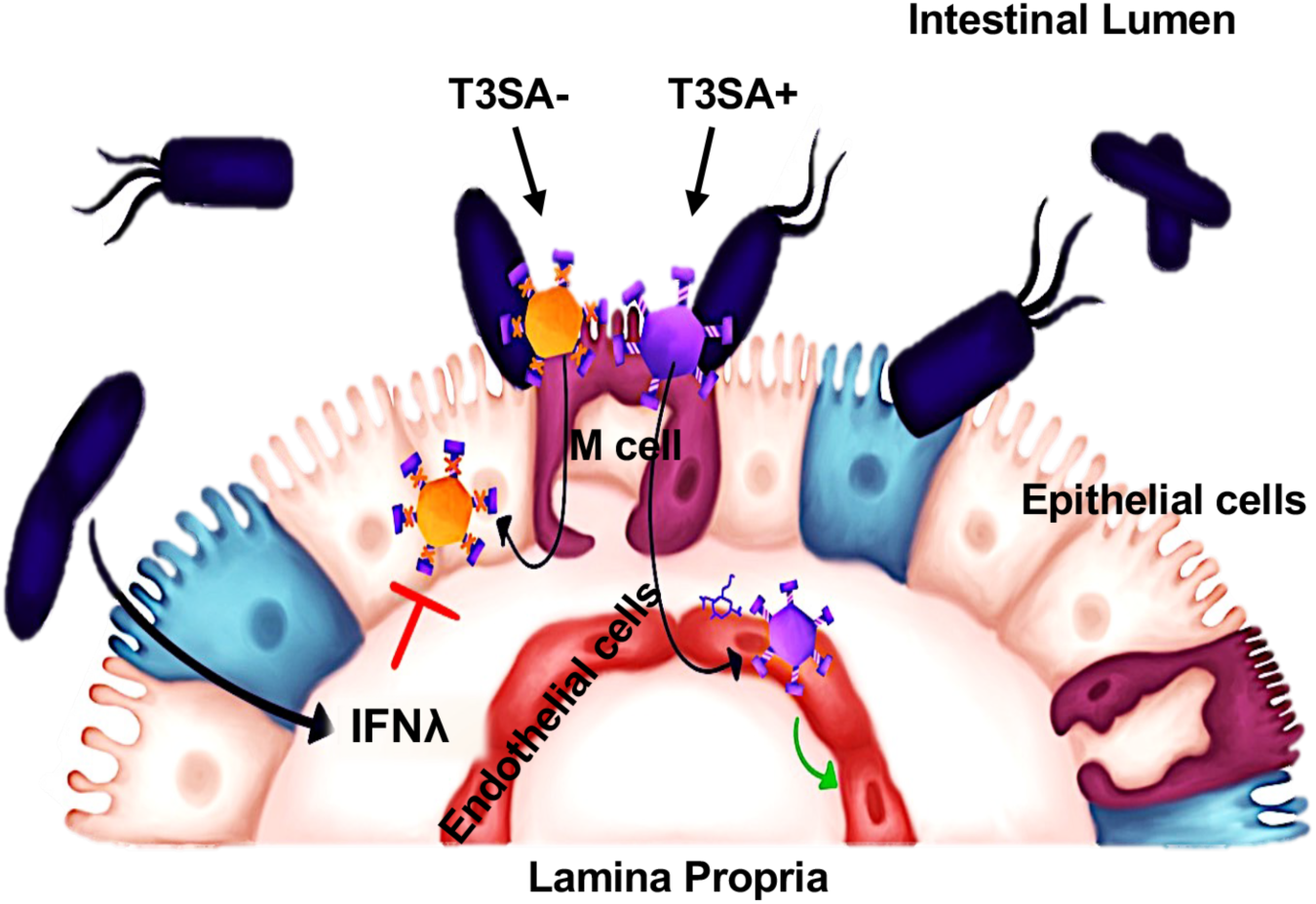
Model for microbiota effects on different reovirus strains. Both T3SA+ (purple) and T3SA− (orange) reovirus strains bind to bacteria (blue) in the intestinal lumen, which may aid delivery of virions through M cells to the underlying lamina propria. Due to its capacity to bind cell surface SA, T3SA+ efficiently infects endothelial cells within the lamina propria while T3SA− infects intestinal epithelial cells from the basolateral side. Microbiota stimulate IFNl responses that limit replication of T3SA− in epithelial cells, but do not affect replication of T3SA+ in endothelial cells. Overall, microbiota promote replication of T3SA+, but limit replication of T3SA−.

## ACKNOWLEDGEMENTS

We are grateful to members of the Dermody and Pfeiffer laboratories for many useful suggestions during the conduct of this research. We thank Megan Baldridge for *Ifnlr1^−/−^* mice, Alexandra Wells for helpful discussions, Cassie Behrendt for training and expertise with germ-free mouse experiments, Melanie Dietrich and Thilo Stehle for sharing the σ1 model, and Kiwi Dammond for artwork. We thank the UTSW Histopathology Core for tissue processing and the Quantitative Light Microscopy Core, a shared resource of the Harold C. Simmons Cancer Center, supported in part by NCI Cancer Center Support Grant P30 CA142543 and S10 OD028630. This work was supported by NIH grants R37 AI074668 and R01 AI158351 to J.K.P., T32 AI005284 to R.W.M., and R01 AI174526 to T.S.D.

## AUTHOR CONTRIBUTIONS

Conceptualization, A.K.E. and J.K.P.; Methodology and Investigation, A.K.E., D.M.S., O.L.W., R.W.M.; Writing – Original Draft, A.K.E. and J.K.P.; Writing – Review and Editing, A.K.E., D.M.S., O.L.W., R.W.M., T.S.D., J.K.P.; Funding Acquisition, T.S.D. and J.K.P.; Supervision, T.S.D. and J.K.P.

## DECLARATION OF INTERESTS

The authors declare no competing interests.

## Key resources table

| REAGENT or RESOURCE | SOURCE | IDENTIFIER |
| --- | --- | --- |
| **Antibodies** |  |  |
| Rabbit anti-reovirus polyclonal antibody | T. Dermody Lab | N/A |
| Donkey Anti-Rabbit IgG H&L (Alexa Fluor® 488) | Abcam | ab150073 |
| Jackson Immuno Research Labs Alexa Fluor 594 AffiniPure Goat Anti-Rabbit IgG (H+L) | Fisher Scientific | NC0414256 |
| Goat anti-Mouse IgG (H+L) Highly Cross-Adsorbed Secondary Antibody, Alexa Fluor™ 594 | Thermo Fisher | A-11032 |
| Rabbit polyclonal Anti-CD45 antibody | Abcam | ab10558 |
| Anti-CD31 monoclonal antibody | Invitrogen | MA5-13188 |
| DAPI Nuclear Counterstain | Thermo Fisher | 62248 |
| **Bacterial and virus strains** |  |  |
| Reovirus T1L | T. Dermody Lab | N/A |
| Reovirus T3SA+ | T. Dermody Lab | N/A |
| Reovirus T3SA- | T. Dermody Lab | N/A |
| Reovirus T3SA- (N198D) | T. Dermody Lab | N/A |
| Reovirus T3SA- (R202W) | T. Dermody Lab | N/A |
| *Bacillus cereus* | Erickson et al., 2018 | N/A |
| *Escherichia coli* K-12 | Erickson et al., 2018 | N/A |
| **Biological samples** |  |  |
| **Chemicals, peptides, and recombinant proteins** |  |  |
| N-Acetylneuraminic Acid, Synthetic - CAS 131-48-6 | Calbiochem, Millipore Sigma | 110138-1GM |
| Neuraminidase (Sialidase) from *Clostridium perfringens* | Sigma Millipore/Roche | 11585886001 |
| NHS-Fluorescein | Thermo Scientific | 46409 |
| **Critical commercial assays** |  |  |
| Molecular Instruments HCR RNA-FISH | https://www.molecularinstruments.com/ | N/A |
| **Deposited data** |  |  |
| **Experimental models: Cell lines** |  |  |
| GFP-Expressing C57BL/6 Mouse Primary Small Intestinal Microvascular Endothelial Cells | Cell Biologics | C57-6054GFP |
| Murine L929 cells | T. Dermody Lab | N/A |
| Murine erythroleukemia cells | T. Dermody Lab | N/A |
| Bovine erythrocytes | Innovative Research | IBORBC100P15ML |
| **Experimental models: Organisms/strains** |  |  |
| Mouse: C57BL/6J | Jackson Labs | Strain #000664 |
| Mouse: C57BL/6J Germ-free | UTSW Gnotobiotics Core | N/A |
| Mouse: C57BL/6J *Ifnlr1*<sup>-/-</sup> | M. Baldrige Lab | N/A |
| **Oligonucleotides** |  |  |
| HCR RNA-FISH Custom Probe Set: Orthoreovirus T1L-S4 | Molecular Instruments | Accession# M13139.1 |
| **Recombinant DNA** |  |  |
| **Software and algorithms** |  |  |
| FIJI | Schindelin et al.<sup>75</sup> | N/A |
| GraphPad Prism V10.2.3 for Mac OS | www.graphpad.com/ | N/A |
| FlowJo-OSX64-10.10.0 | www.flowjo.com/ | N/A |

## METHODS

### Viruses and cells

Reovirus strains were propagated, purified, and quantified by plaque assay using L929 cells as previously described^28^. Mutant reovirus strains were engineered using site-directed mutagenesis as previously described^76^. Purified T3SA+ and T3SA− virions were fluorescently-labeled as previously described^16^. Briefly, viral particles (2 × 10^12^, determined by A_260_ of 1 = 2.1 × 10^12^ viral particles/mL) were diluted in sodium bicarbonate (0.05 M, pH 8.5) to a total of 495 μL, and 5 μL of 1 mM NHS-Fluorescein was added to virus samples or controls without virus. Dye was incubated with virus at 4°C for 90 min in the dark, and unbound dye was removed by dialysis. Viral particle numbers (A_260_) and titers of fluorescently-labeled viral stocks were determined as described^16^. Murine L929 cells were maintained in DMEM supplemented to contain 5% FBS, 2 mM L-glutamine, and antibiotics (penicillin/streptomycin). Suspension cultures of murine erythroleukemia (MEL) cells were maintained in F12 medium supplemented to contain 10% FBS, 2 mM L-glutamine, and antibiotics (penicillin/streptomycin).

### Primary cell lines

Primary small intestine microvascular endothelial cells were purchased from Cell Biologics, maintained in complete endothelial cell medium (Cell Biologics, M1168), and propagated at 37°C in 5% CO_2_ until confluent and passaged no more than 2 times prior to experimental use.

### Mice and animal experiments

All animals were handled according to the Guide for the Care of Laboratory Animals of the NIH. All mouse studies were conducted at UT Southwestern (Animal Welfare Assurance #A3472-01) using protocols approved by the local Institutional Animal Care and Use Committee. C57BL/6J wild-type mice were obtained from the Jackson Laboratory, C57BL/6J mice defective for the interferon lambda receptor 1 (*Ifnlr1^−/−^*) were obtained from Megan Baldridge at Washington University in St. Louis^5^. All experimental mice were 3-5 weeks old, with equal numbers of male and female mice, and cages and sex were assigned randomly to groups to limit differences caused by litter-to-litter and microbiota variability. Microbiologically-sterile germ-free C57BL/6J mice were maintained in gnotobiotic chambers until use, and housed in the germ-free BSL2+ facility in sterile cages during experiments. Microbiota colonization of germ-free mice was conducted as previously described^11^. Briefly, seven days prior to reovirus infection, germ-free mice were perorally-administered 25 μl of fecal slurry (2-3 fecal pellets in 500 μl PBS) from untreated conventional mice. For antibiotic treatments, germ-free or conventional C57BL/6J mice were perorally administered an antibiotic cocktail containing vancomycin, ampicillin, neomycin, metronidazole, and streptomycin for 5 days (10 mg each antibiotic in 25-50 μL water per mouse once daily) followed by *ad libitum* antibiotic administration in drinking water (500 mg/L vancomycin, and 1 g/L ampicillin, neomycin, metronidazole, and streptomycin) for the duration of experiments. For exogenous *N*-acetylneuraminic acid/sialic acid (SA) treatments mice were orally administered 1 mg of SA in 10 μL water twice/day for 2 days prior to infection as well as in drinking water at 1% concentration throughout the experiment. Mice were perorally inoculated with 10^8^ PFU of T1L, T3SA+, T3SA−, or 5 × 10^7^ PFU of T3SA− (N198D) or T3SA− (R202W). Feces and intestinal tissues were collected at 4 dpi and processed for viral titer or HCR analysis.

For viral titer assays, the intestine was flushed with PBS, dissected into sections (duodenum, jejunum, ileum, cecum, or colon), allocated into separate tubes, and frozen at −80°C. Tissues were weighed, resuspended in 0.5 g/mL PBS, homogenized using 0.9-mm stainless steel beads in a Bullet Blender Tissue Homogenizer (Next Advanced Inc) for 5 min and frozen and thawed twice. Cell debris was removed by centrifugation at 13,000 RPM for 3 minutes. Virus in the supernatant was quantified by plaque or FFU assay using L929 cells. For HCR-FISH, the entire small intestine was collected, flushed with 10 mL PBS, flushed with 5 mL of ice-cold 4% paraformaldehyde/DEPC-PBS (fixative), and incubated in 20 mL of 10% neutral-buffered formalin at 4°C for 2-4 h. Small intestine was then Swiss-rolled, paraffin-embedded, cut into 5 μm sections per slide (UTSW Histo Pathology Core).

### HCR-FISH and quantification of reovirus-infected cells in the intestine

HCR-FISH was conducted using reagents purchased from Molecular Instruments and according to the HCR™ RNA-FISH (v3.0) or HCR™ IF + HCR™ RNA-FISH protocols for formalin-fixed paraffin-embedded human tissue sections^58,77^. Slides were deparaffinized using xylene and rehydrated with decreasing concentrations of ethanol washes. Antigen unmasking was conducted using citrate buffer. For protein detection, tissues were incubated with primary antibody, CD45 (1:1000) or CD31 (1:100), in antibody buffer at 4°C overnight, washed and incubated with 1 μg/mL of initiator-labeled secondary (CD45: Donkey Anti-Rat-647; CD31: Donkey Anti-Mouse-594, Molecular Instruments) at room temperature for 1 h.

Reovirus T3SA+ or T3SA− viral RNA (vRNA) was detected using custom probes designed by Molecular Instruments (Accession# M13139.1) to specifically target the reovirus T1L S4 genome segment. Tissues were incubated with 100 μL of probe solution containing 30 nM reovirus S4 probes and incubated at 37°C overnight. Tissues were washed according to manufacturer’s instructions, and 488-labeled specific hairpins added in amplification buffer and incubated at room temperature for < 16 h. Fluorescent hairpins were thoroughly washed off, tissues stained with DAPI (1:1000) prior to mounting with ProLong Diamond (Invitrogen). Slides were cured overnight and imaged using a Nikon SoRa spinning-disk confocal microscope. For quantification of reovirus vRNA localization within the intestinal villi, 20-40X magnification was used and 3-5 different areas per slide were imaged, capturing ~15-40 villi per image. Images were then used to count vRNA-positive foci localized within the IEC and/or LP of 100 total villi. To compare IEC or LP cellular localization of T3SA+ and T3SA− across treatment conditions, we normalized the data to a set control condition: the average total (IEC+LP) number of T3SA+ infected foci in wild-type, conventional mice. Specifically, for each condition, the mean was calculated as the number of positive IEC or LP foci divided by the average number of total positive foci (IEC+LP) in control/conventional WT mice infected with T3SA+, which was 688 reovirus-positive foci from 500 villi in 5 mice giving an average of (688÷5) 138 [Mean = [(# foci in LP or IEC) ÷ (control, 138)] × 100%].

### Reovirus replication and Fluorescent Focus Unit (FFU) assays

Viral replication efficiency was determined following infection of murine L929 cells and murine erythroleukemia (MEL) cells as previously described^78^. L929 cells were seeded into 24-well plates, incubated overnight, and adsorbed with reovirus at an MOI of 0.5 at room temperature for 1 h. Inoculum was removed, and cells were incubated in 1 mL of fresh medium at 37°C. At 0, 6, 12, 18, or 24 h post-adsorption, plates were frozen and thawed twice to release intracellular virus, and viral titers determined by plaque assay using L929 cells. For reovirus replication in MEL cells, 5 × 10^4^ cells were suspended in 100 μL culture medium containing reovirus at an MOI of 2 and incubated with gentle agitation at room temperature for 1 h. Cells were collected by centrifugation, resuspended in 1 mL of fresh medium, and transferred to 24-well tissue-culture plates. At 0, 6, 12, 18, or 24 h post-adsorption, cells were frozen and thawed twice prior to determination of viral titer by plaque assay using L929 cells.

For quantification of reovirus infected cells by FFU assay, L929 or mouse primary small intestinal microvascular endothelial cells were plated in 24-well plates and maintained at 37°C in 5% CO_2_ until semi-confluent or 10^5^ cells per well. For removal of cell surface SA, cells were incubated with 80 mU/mL of neuraminidase in PBS at 37°C for 1 h. For treatment with excess SA, 10^4^ PFU (determined by plaque assay using L929 cells) of T3SA+ or T3SA− was incubated with 10 mg/mL *N*-Acetylneuraminic Acid (SA) in PBS at room temperature for 1 h. Cells in triplicate wells were absorbed with reovirus T3SA+ or T3SA− at an MOI of 0.01 at room temperature for 30 min. Inoculum was removed and culture medium added to cells. At 18 hpi, cells were fixed with ice-cold methanol and fluorescent focus assays were conducted to detect reovirus-infected cells. The average of triplicate wells was quantified as fluorescent focus units (FFU)/mL. Reovirus FFU assays were conducted using rabbit polyclonal reovirus-specific antiserum (1:10,000 dilution in 1%BSA/PBS) and Alexa Fluor 594 or 488-conjugated anti-rabbit secondary antibody (1:500 dilution in 1%BSA/PBS). Nuclei were stained with DAPI. Cells were imaged using Nikon fluorescence microscope and quantification of reovirus infected foci determined manually or with FIJI software.

### Hemagglutination assay

Hemagglutination assays were conducted as previously described^78^. Briefly, purified reovirus particles (10^11^) were serially diluted two-fold from 5×10^10^ to 2.4×10^7^ viral particles in PBS (50 μL) within 96-well U-bottom microtiter plates (Costar). Bovine erythrocytes were washed, diluted to 1% in PBS, and equal volume added to virus solutions for 100 μL total volume per well. Plates were incubated at 4°C for 3 h and hemagglutination (HA) titers were determined as the reciprocal of the dilution fraction in which the minimum number of particles produces a complete shield of erythrocytes.

### Bacterial binding assays

*Escherichia coli* and *Bacillus cereus* are lab strains and previously described^15^. For binding assays, frozen glycerol stocks were used to inoculate 5 ml liquid cultures in brain heart infusion medium, incubated at 37°C overnight with shaking, and the following morning, subcultures were inoculated and grown for 4-6 h, centrifuged, washed, and pellets resuspended in 1 mL PBS. Bacterial concentrations were determined by OD_600_ measurements and diluted to 10^7^ CFU/mL and incubated with NHS-Fluorescein-mock, or 10^10^ NHS-Fluorescein-labeled T3SA+ or T3SA− viral particles at room temperature for 15 min with rotation. Bacteria were centrifuged at 8,000 RPM for 5 min, unbound virus was removed, and cells were gently washed and resuspended in 100 μL 1%PFA/1%BSA/PBS. Bacteria were stained with DAPI (1:1000) for 5 min, diluted to 1 mL in PBS, and 10^4^ CFU bacteria were added to microscope slides. Slides were dried and mounted for confocal microscopy and imaged using a Nikon SoRA spinning-disk confocal microscope at 60X magnification. For flow cytometry assays, bacterial binding was conducted as described above, but bacteria were incubated with PBS or unlabeled purified T3SA+ or T3SA− stocks. After binding, bacteria were stained with reovirus-specific polyclonal serum (1:10,000), Alexa Fluor 488-conjugated anti-rabbit secondary antibody (1:500), and DAPI. Bacteria were washed to remove unbound antibodies and resuspended to 10^6^ CFU in 100 μL of 1%PFA/1%BSA/PBS and analyzed using a FACSCalibur cytometer (Becton Dickinson, Franklin Lakes, NJ). The percentage of bacterial cells bound by reovirus (488+) was determined from 10^5^ total live bacteria counts using FlowJo software.

## QUANTIFICATION AND STATISTICAL ANALYSIS

Data were analyzed using GraphPad Prism and are presented as the mean values ± standard error of mean (SEM). The figure legends report the number of biological replicates (n), the number of independent experiments, and the statistical tests conducted. Differences were analyzed using either multiple unpaired t tests, ordinary one-way ANOVA with Fishers LSD test, ordinary one-way ANOVA and Dunnett’s test, or Brown-Forsythe and Welch ANOVA tests. Differences were considered significant at *P* < 0.05.

## SUPPLEMENTAL FIGURE AND TABLE LEGENDS

**Supplemental Figure 1.**
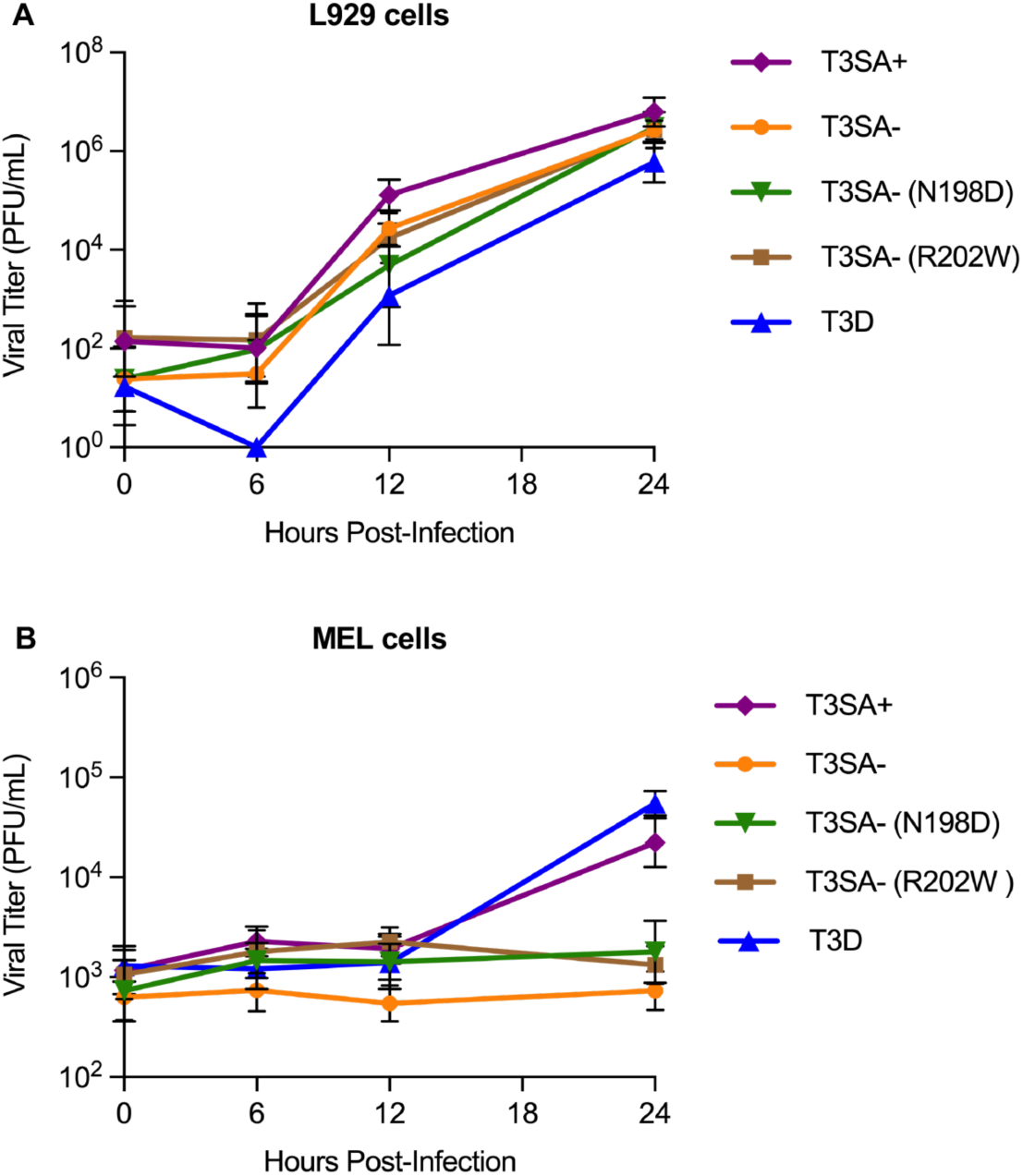
Reovirus mutants incapable of binding SA can replicate in highly susceptible L929 cells but not in murine erythroleukemia (MEL) cells, which require viral SA binding for infection. (A,B) L929 cells (A) or MEL cells (B) were adsorbed with T3D, T3SA+, T3SA−, or mutant strains T3SA−/N198D or T3SA−/R202W at a multiplicity of infection (MOI) of 0.5 (A) or 2 (B). Titers of virus in cell lysates at the times post-adsorption shown were determined by plaque assay using L929 cells. Results are shown as mean ± SEM (n=6 per time point) from three independent experiments.

**Supplemental Table 1.** Distinct cell tropisms of reovirus strains T3SA+ and T3SA− in the small intestines of conventional or antibiotic-treated C57BL/6 (WT) and *Ifnlr1^−/−^* mice. C57BL/6 (WT) or *Ifnlr1^−/−^* (cyan) mice were treated with or without antibiotics prior to peroral inoculation with 10^8^ PFU of T3SA+ (purple outline) or T3SA− (orange outline). At 4 dpi the small intestine was harvested, Swiss rolled, and processed for HCR-FISH analyses for reovirus-infected foci (see Figure 4A legend). The total numbers of reovirus-infected intestinal epithelial cells (IEC) or lamina propria (LP) foci in 100 intestinal villi per mouse were determined from confocal images. The data are presented as the mean ± SEM from 5 WT or 3 *Ifnlr1^−/−^* mice per condition relative to the total positive-foci in WT conventional mice infected with T3SA+, which was set to 100% (with 78% of foci in lamina propria cells and 22% of foci in epithelial cells). For each condition, the mean was calculated as the number of positive epithelial foci or lamina propria foci divided by the average number of positive foci in conventional WT mice infected with T3SA+, which was 138 foci from 500 total villi in 5 mice, multiplied by 100%. Means are overlaid with heat maps to highlight lowest to highest values demarcated as shown in the color scale. *P*-values were assessed using an unpaired t test between the indicated groups.

| Intestinal Epithelial Cell (IEC) |  |  |  |  |  |  |  |
| --- | --- | --- | --- | --- | --- | --- | --- |
| Virus | Mouse | n | ABX | Mean | SEM | Conv vs ABX | WT vs *Ifnlr1*<sup>-/-</sup> |
|  |  |  |  |  |  | *P*-value | *P*-value |
| T3SA+ | WT | 5 | - | 22 | 4.9 | 0.030 | 0.100 |
| T3SA+ | WT | 5 | + | 9 | 1.5 |  |  |
| T3SA+ | *Ifnlr1*<sup>-/-</sup> | 3 | - | 39 | 7.4 | 0.018 | 0.646 |
| T3SA+ | *Ifnlr1*<sup>-/-</sup> | 3 | + | 8 | 2.9 |  |  |
| T3SA- | WT | 5 | - | 10 | 2.8 | 3E-05 | 0.001 |
| T3SA- | WT | 5 | + | 41 | 2.1 |  |  |
| T3SA- | *Ifnlr1*<sup>-/-</sup> | 3 | - | 45 | 6.6 | 0.702 | 0.229 |
| T3SA- | *Ifnlr1*<sup>-/-</sup> | 3 | + | 49 | 7.4 |  |  |
| Lamina Propria (LP) |  |  |  |  |  |  |  |
| Virus | Mouse | n | ABX | Mean | SEM | Conv vs ABX | WT vs *Ifnlr1*<sup>-/-</sup> |
|  |  |  |  |  |  | *P*-value | *P*-value |
| T3SA+ | WT | 5 | - | 78 | 11.6 | 0.001 | 0.688 |
| T3SA+ | WT | 5 | + | 18 | 3.3 |  |  |
| T3SA+ | *Ifnlr1*<sup>-/-</sup> | 3 | - | 69 | 16.2 | 0.035 | 0.969 |
| T3SA+ | *Ifnlr1*<sup>-/-</sup> | 3 | + | 18 | 2.6 |  |  |
| T3SA- | WT | 5 | - | 2 | 0.4 | 0.018 | 0.005 |
| T3SA- | WT | 5 | + | 5 | 1.1 |  |  |
| T3SA- | *Ifnlr1*<sup>-/-</sup> | 3 | - | 8 | 2.0 | 0.346 | 0.020 |
| T3SA- | *Ifnlr1*<sup>-/-</sup> | 3 | + | 11 | 1.5 |  |  |

Color Scale

## Notes

### Competing Interest Statement

The authors have declared no competing interest.

### Summary of Updates

Corrected spelling of author name.

